# The influence of major S protein mutations of SARS-CoV-2 on the potential B cell epitopes

**DOI:** 10.1101/2020.08.24.264895

**Authors:** Xianlin Yuan, liangping li

**Affiliations:** Department of Oncology and Institute of Clinical Oncology, The first Affiliated Hospital, Jina University, Guangzhou, Guangdong, People’s Republic of China; Department of Immunology, Zhongshan School of Medicine, Sun Yat-sen University, Guangzhou, Guangdong, People’s Republic of China

**Keywords:** SARS-CoV-2, Spike protein, Receptor-binding domain, Mutation, Variant, Neutralizing antibody, Vaccine

## Abstract

SARS-CoV-2 has rapidly transmitted worldwide and results in the COVID-19 pandemic. Spike glycoprotein on surface is a key factor of viral transmission, and has appeared a lot of variants due to gene mutations, which may influence the viral antigenicity and vaccine efficacy. Here, we used bioinformatic tools to analyze B-cell epitopes of prototype S protein and its 9 common variants. 12 potential linear and 53 discontinuous epitopes of B-cells were predicted from the S protein prototype. Importantly, by comparing the epitope alterations between prototype and variants, we demonstrate that B-cell epitopes and antigenicity of 9 variants appear significantly different alterations. The dominant D614G variant impacts the potential epitope least, only with moderately elevated antigenicity, while the epitopes and antigenicity of some mutants(V483A, V367F, etc.) with small incidence in the population change greatly. These results suggest that the currently developed vaccines should be valid for a majority of SARS-CoV-2 infectors. This study provides a scientific basis for large-scale application of SARS-CoV-2 vaccines and for taking precautions against the probable appearance of antigen escape induced by genetic variation after vaccination.

**Author Summary:** The global pandemic of SARS-CoV-2 has lasted for more than half a year and has not yet been contained. Until now there is no effective treatment for SARS-CoV-2 caused disease (COVID-19). Successful vaccine development seems to be the only hope. However, this novel coronavirus belongs to the RNA virus, there is a high mutation rate in the genome, and these mutations often locate on the Spike proteins of virus, the gripper of the virus entering the cells. Vaccination induce the generation of antibodies, which block Spike protein. However, the Spike protein variants may change the recognition and binding of antibodies and make the vaccine ineffective. In this study, we predict neutralizing antibody recognition sites (B cell epitopes) of the prototype S protein of SARS-COV2, along with several common variants using bioinformatics tools. We discovered the variability in antigenicity among the mutants, for instance, in the more widespread D614G variant the change of epitope was least affected, only with slight increase of antigenicity. However, the antigenic epitopes of some mutants change greatly. These results could be of potential importance for future vaccine design and application against SARS-CoV2 variants.

## Introduction

A severe contagious pneumonia caused by a novel coronavirus was first reported outbreaking at Wuhan of China in December 31, 2019 and soon detected in other countries during few months [1]. This disease was formally named as coronavirus disease 2019 (COVID-19) by World Health Organization (WHO). The genome sequence of the pathogen was soon identified by NGS (accession number: QHO62107.1 from NCBI database) to be a novel beta coronavirus, belonging to the family coronaviradae. As its genome and pathology of the disease is similar to severe acute respiratory syndrome coronavirus (SARS-CoV) breaking out in 2009 [2], this virus was named as SARS-CoV-2. In view of the phylogenetic analysis [1], SARS-CoV-2 shares 79.6% sequence identity with SARS-CoV [3] and 50% with Middle-East respiratory syndrome coronavirus (MERS-CoV) [4]. Although similar in genome, SARS-CoV-2 is far more contagious, much faster spreading and more destructive than other types of coronavirus: SARS-CoV [5] and (MERS-CoV) [6]. Since January 30, 2020, the WHO announced the CoVID-19 contagion as a public health emergency of global concern. As of July 20, 14348858 cases of COVID-19 and 603691 deaths have been reported globally according to COVID-19 Situation Report–182 (WHO website at https://www.who.int/emergencies/diseases/novel-coronavirus-2019).

The virion of SARS-CoV-2 is spherical, enveloped, and 60-140 nm in diameter with spikes of about 9-12 nm outside. The coronaviral genome encodes 10 proteins, four of them are major structural proteins: the spike (S), membrane (M), envelope (E) and nucleocapsid (N) proteins [7]. Each of these proteins is responsible for different functions in the life cycle of the virus: M protein decides the shape and pattern of the virus envelope. The viral assembly and germination was accomplished by E protein. N proteins and RNA genome of virion are closely linked and participates in viral replication and assembly. Most importantly, the S protein is like a bridge to attach and bind to the host cell receptors, and results in the fusion of the viral and host cellular membranes and consequential viral entry into host cell [7]. The S protein, a I-type transmembrane glycoprotein, compose of ectodomain, transmembrane domain (TM) and CT domain. The ectodomain is made up of two subunits (S1 and S2): the S1 subunit includes N-terminal domain (NTD) and receptor-binding domain (RBD), and S2 subunit contains fusion peptide (FP), heptad repeat (HR) domain 1 and 2. The RBD domain is responsible for binding to the receptor of host cells angiotensin-converting enzyme 2 (ACE2), while S2 subunit completes the mission of viral fusion and entry [8]. Previous studies on SARS-CoV had shown that RBD was major targets of effective neutralizing antibodies. Therefore, S protein not only is a trigger of virus replication and transmission, but also is the key target of the SARS-CoV-2 vaccine for prevention of Covid-2019.

However, owing to the extensive transmission of SARS-CoV-2, the genetic variants of the virus have appeared in a growing number of countries. 5775 mutations in the SARS-CoV-2 genome were discovered from 10022 public genome data assemblies as at May 1, 2020 [9], in which 394 missense mutations of S protein were detected. Among these spike mutations, D614G mutation, in which Aspartic acid (D) was replaced with Glycine (G) at the AA site of 614, was a major mutation of great concern [10, 11]. SARS-CoV-2 with D614G mutation may have triggered fatal infections in many European countries, such as Spain, Italy, France, etc. [11].

These mutations will undoubtedly cause changes in the structure of S proteins. However, it’s highly worth concerning whether or not these mutations affect the antigenicity of S proteins and the binding ability with neutralizing antibodies. If the B-cell epitopes on S protein changed and could not bind the neutralizing antibodies, it would result in losing efficacy of the developed vaccines based on prototype S protein.

Many immuno-bioinformatic tools have been developed to dope out the overall and deep analysis of viral antigens, including both linear and discontinuous epitopes of B-cells as well as their immunogenicity, etc. To explore these questions, here we report to used these immuno-bioinformatic tools from the IEDB and related resources to predict the B cell epitopes of S protein from the prototype and mutated strains of SARS-CoV-2 and compare the changes of the likely epitope sites from dominant and rare mutations of S protein. We found that the distinctive mutations of S proteins could impact potential effective epitopes of S proteins in different degree.

## Results

### Epidemiology statistics of S protein mutations

We first searched and compared the epidemiology statistic data of S protein mutations published by Korber B and Takahiko Koyama. Korber B et al collected 4,535 genome sequences as of April 13^th^; Koyama et al identified 5775 distinct genome variants from 10 022 SARS CoV-2 genomes, which submitted into database before May 1, 2020 [12]. In the Koyama’s report, S protein contained 394 missense mutations [9]. In both reports, the mutation D614G was of the highest frequency and other mutations were relatively rare. Through quantitative statistics of major variants at two time points, we found that in just two weeks, the number of D614G mutation was doubled, which demonstrated that this variant was dominant form among S protein mutations. The frequency of other mutations changed slightly, and some showed zero growth (**Table 1**).

### Conservation analysis of the prototype S protein

We chose the earliest submitted strain (NCBI ID:QHO62107.1) detected at China as prototype of S protein. Before predicting the B-cell epitopes, we carried out the conservation analysis by comparing this sequence with other earlier sequence submitted from different countries at January-February, 2020. Sequence of SARS-CoV2 S protein from 10 different countries isolates including China (NCBI ID:QHO62107.1), Japan (BBW89517.1), USA (QHQ82464.1), Germany (QKM76570.1), Egypt (QKS66892.1), Spain (QKJ68388.1), France (QJT72638.1), Greece (QIZ16535.1), Australian (QHR84449.1) and Russia (QKV28206.1) were subjected to multiple sequence-alignment through Clustal Omega tool (Figure S1). Conservation analysis of S protein sequence manifests that the prototype have 100% identity with all the retrieved sequences. Thus, we used the earliest version of S protein sequence as prototype for following studies of B-cell epitope prediction.

### Structural analysis of prototype S protein

To localize these mutations in the position of different functional domains of S protein, we analyzed S protein sequence by bioinformatics tools. First to analyze the trans-membrane protein topology, we applied the online tool TMHMM to treat the sequence of S protein, and localized one transmembrane region. The spatial distribution of residues could be divided into three parts: residues from 1 to 1213 on the extracellular surface, residues from 1214 to 1236 in the region of transmembrane (TM) and residues from 1237 to 1273 in the cytoplasmic region (CT).

The exracellular domains divide into S1 and S2 subunits; S1 contains the N-terminal domain (NTD) and receptor binding domain (RBD) [13]. Based on the beginning and end position of different domains, 10 major mutations were shown in the schematic diagram of S protein (Figure 1A). We found that most of S protein mutations (70%) locates in S1 and near region of S protein. The highest mutation D614G is near RBD domain.

**Fig 1.**
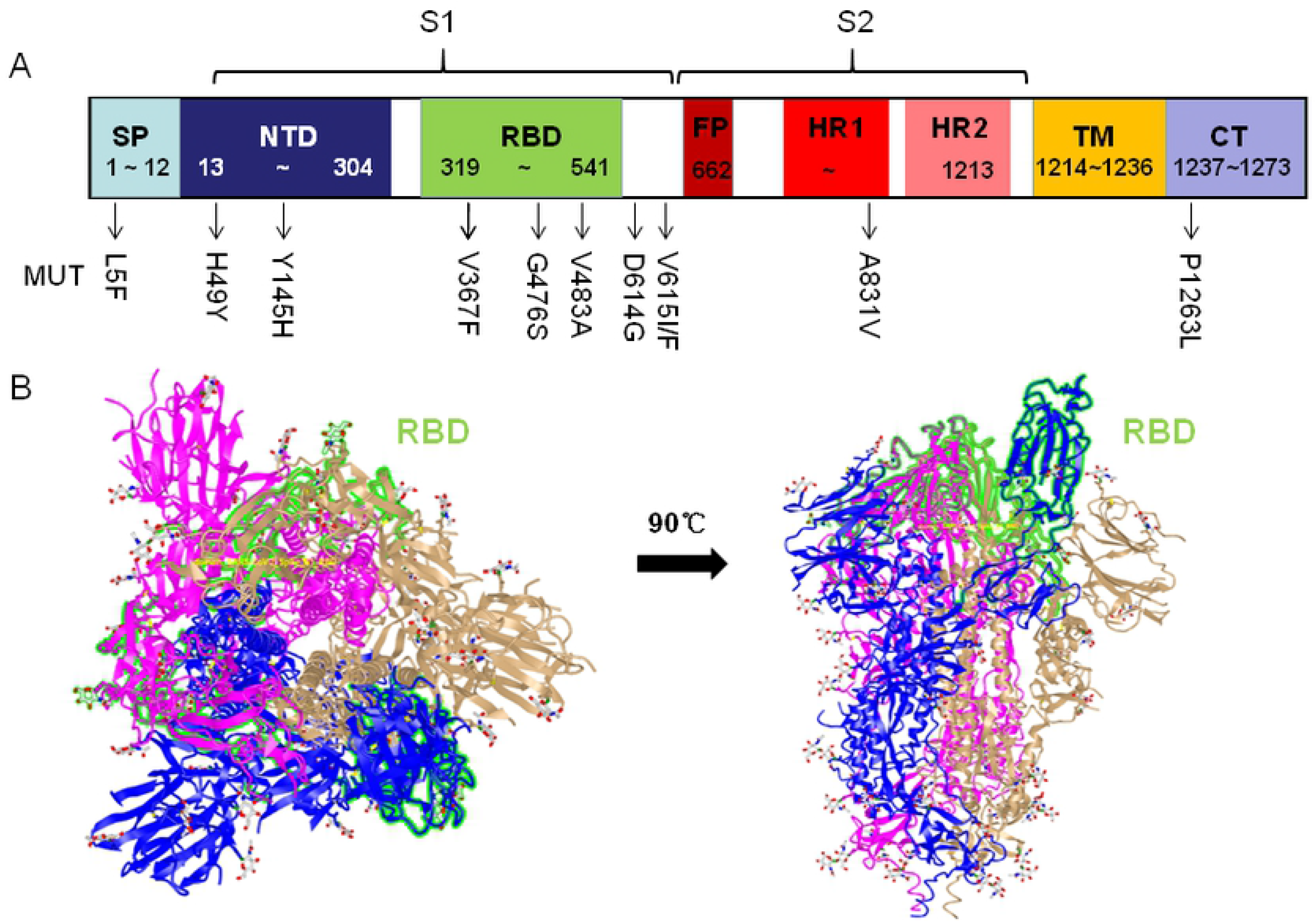
Structural domains and mutation positions of SARS-COV-2 S protein. (A) The schematic diagram of structural domains and mutation positions of SARS-COV-2 S protein. 8 domains of S protein are labeled with different colored box. The major common mutations and positions are shown under the domain boxs. SP, signal peptide; NTD, N-terminal domain; RBD, receptor binding domain; FP, fusion peptide; HR1, heptad repeat 1; HR2, heptad repeat 2; TM, transmembrane region; CT, cytoplasmic region. (B) 3D structure of S protein (6vyb, quoted from Walls, A.C etc.). The 3D structure of S protein is displayed in two directions. RBD domain indicated by green highlight is shown in iCn3D Viewer.

To show the functional domains at a 3D level and analyze disc-continuous epitopes of S protein, we search database through homology modeling of swiss-model tool, and found out the 3D structure file of S protein PDB ID: 6vyb, in which the amino acid sequences were 99.5% consistent with spike glycoprotein. SARS-CoV-2 spike protein is a trimer, and the 3D structure of ectodomain (open state) from 6vyb is shown in Figure 1B [14], which contains three chains of A/B/C.

### Prediction of B-cell epitopes on prototype S protein

Humoral immunity play a very important role in defense of viral infection. The B cell receptor (BCR) or neutralizing antibodies recognize B cell epitopes (linear and discontinuous) of S protein, which generally exist on the virus surface as natural antigen molecules without processing.

To predict the potential linear B-cell epitopes, we first used BepiPred-2.0 prediction tool on IEDB server to screen the prototype S protein sequence and discovered total 30 B-cell linear epitopes (Table S2), whose distribution is shown on Figure 2A. Most of the B-cell epitopes are located on the NTD and RBD domains of S protein. Next, we determined the effective epitopes by analyzing the antigenicity with Vaxijen 2.0 tool and accessibility with Emini Surface Accessibility Prediction tool (Figure 2B). Totally, 12 effective epitopes were found from 30 predicted epitopes. The position, sequence, length and evaluation scores of potential B-cell linear epitopes are listed in **Table 2**. Among them, 9 epitopes are in S1 subunit 4 in the NTD region, 5 in the RBD domain) and 3 in the S2 subunit of S protein. Based on this analysis, we found that three epitopes in the RBD domain (_384_PTKLNDL_390_, _405_DEVRQIAPGQTGKI_418_, and _487_NCYFPL_492_) have more significant antigenicity and accessibility.

**Fig 2.**
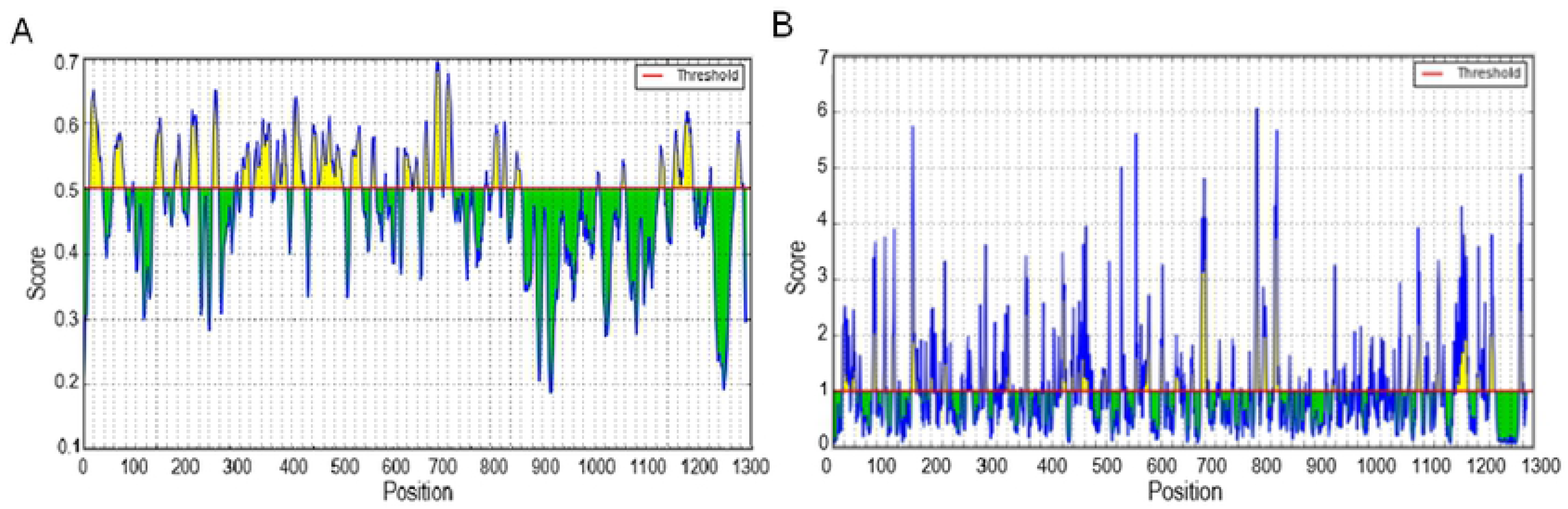
Prediction of B cell linear epitopes and accessibility analysis of prototype S protein. (A) The distribution of all the predicted B-cell linear epitopes by BepiPred-2.0. The residues with scores above the threshold (value is adjusted at 0.55) are predicted to be potential epitopes and colored in yellow. Y-axes indicates residue scores and X-axes exhibits residue positions of the S protein. (B) The surface accessibility analyses using Emini surface accessibility scale. The residues with scores above the threshold (the default value is 1.00) are predicted to have good accessibility.

We further predicted the discontinuous epitopes by the Discotope 2.0 online server. 3D structure of S protein (PDB ID: 6vyb, Chain ID: A) was utilized to predict the discontinuous epitopes. The default threshold was −3.7 with 47% of Sensitivity and 75% of Specificity. The 53 discontinuous epitopes were predicted and mainly located in the whole RBD region at 400aa~600aa of S protein shown in Figure 3A. All of the predicted epitopes distributing on surface of S protein are shown in a 3D structure picture in Figure 3B using JSmol Viewer. According to the distribution in different domains, these epitopes (Table S3) could be divided into four groups (Table 3) and the highest propensity score (P-Score) and DiscoTope score (D-Score) of epitopes were concentrated at 498~500aa of RBD region shown by arrows in the Figure 3B.

**Fig 3.**
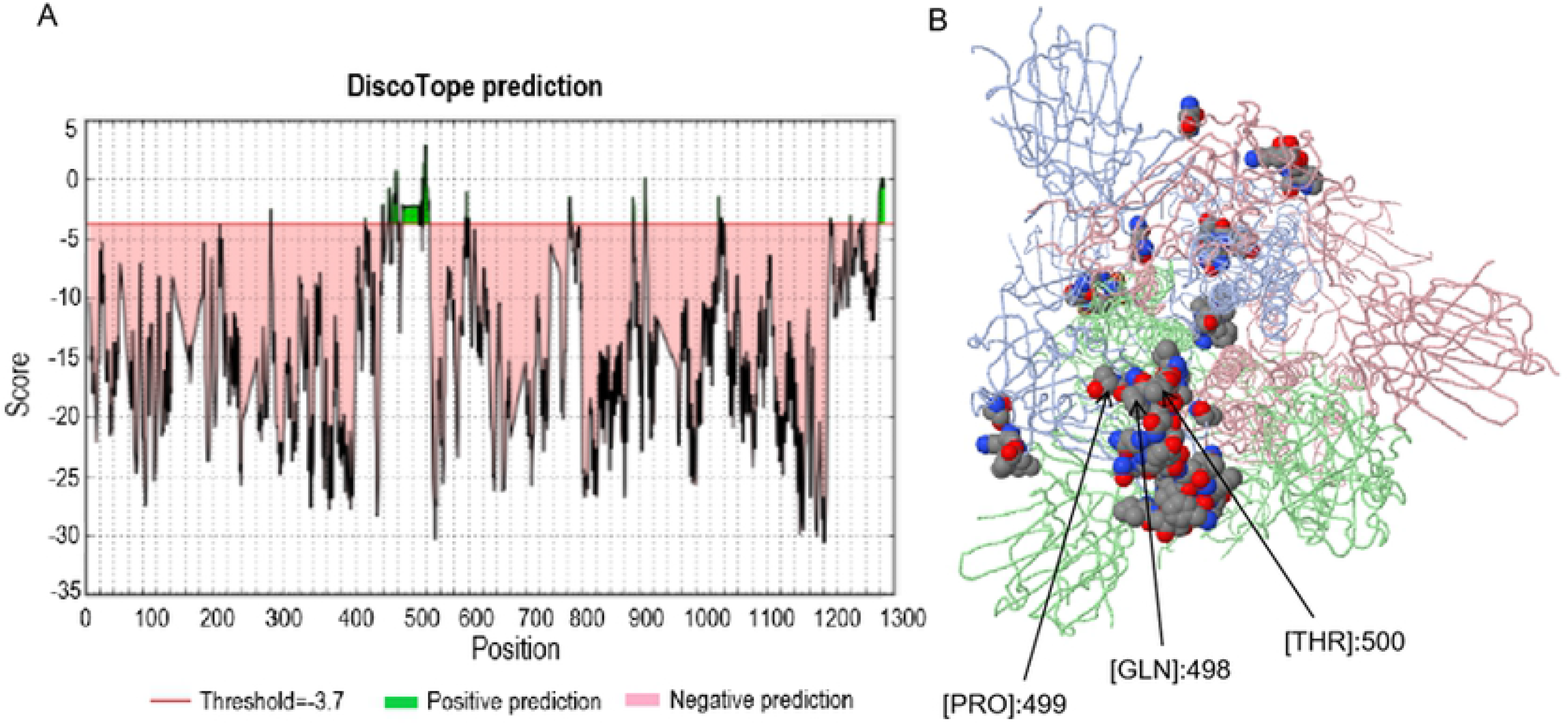
Prediction and distribution of B cell discontinuous epitopes of prototype S protein. The discontinuous epitopes of S protein were predicted by the Discotope 2.0 tool and the default threshold was −3.7. (A) The distribution of the predicted discontinuous epitopes in prototype S proteins. The green part in the figure represents the possible presence of discontinuous epitopes from RBD domain, while the pink region indicates those unlikely epitopes. (B) The surface position of discontinuous antigen epitopes in the 3D structure of prototype S proteins. The arrows in the figure refer to residues from RBD domain with the highest DiscoTope score, which could have the potential to induce a better immune response.

Finally, these epitopes were validated by Pepitope tool (http://pepitope.tau.ac.il/), the three major antigen clusters were consistent with B-linear epitopes mentioned above (Table 4), which further indicated the rationality of our predicted B-linear epitopes.

### Prediction of B-cell epitopes of major variant S protein

The mutations of S protein may influence its structure and change B-cell epitopes for the neutralizing antibody. The missense mutations will result in changes of amino acid residues and may affect B cell epitopes. To assess the impact of the dominant and rare mutations of S-protein to linear B-cell epitopes, we selected those S protein mutations more than 10 counts from these 394 missense mutations to predict B-cell epitopes. Among the several common mutations, we focus the analysis of B-cell epitopes on the ectodomains of S protein, except the L5F mutation in the signal peptide and P1263L mutation in the intracellular region, which are impossible to appear on the surface of the virus. Due to lack of 3D structure data of mutated S proteins, we could not predicted their discontinuous epitopes of B-cells. Through analyzing the sites of 9 missense mutation, we found that these conformational epitopes do not contain any sites of common mutations, which can be inferred that the mutation of S protein has a slight effect on the conformational epitope of B cells.Thus, we predicted the linear B cell epitope of 9 variants as described in following and further determined the changes of epitopes by comparison with the epitopes from the prototype S protein.

#### H49Y mutation

H49Y mutation occurred mainly in China, but seem to be reducing in overall frequency at the present stage [12]. By assessment for its antigenicity and surface availability, we found that four epitopes have changed (Table S4). In brief, after H49Y mutation, the S protein had 14 effective epitopes, two of which have better antigenicity than original epitopes at site of 405~417 and 697~709, two of which were newly generated at sites of 519~533 and 618~629, and the remaining 10 epitopes were the same as those without mutation.

#### Y145H mutation

Y145H mutation occurred in 8 countries, but the frequency appeared to decrease now [12]. By using the screening methods above, we found that five altering sites have distinct influences on the likely epitopes (Table S5). Y145H mutation emerging, the S protein had 13 effective epitopes, two of which were newly produced from originally unlikely epitopes at sites of 618~625, three of which have better antigenicity than original epitopes at site of 140~153, 459~465 and 657~663, and the remaining 9 epitopes were conservative.

#### V367F mutation

V367F mutation existed in Europe and Hong Kong and appeared to be declining in overall global distribution [12]. Only 6 alterations had obvious effects on likely epitopes (Table S6): two previously effective epitopes (208~220 and 487~492) were out of work in replacement for 210~221 and 487~489, one epitope was deleted at site of 459~464, two alterations (141~152→140~154 and 208~220→210~221) greatly reduced the antigenicity of the original epitopes. However, the antigenicity of two previous epitopes (385~392 and 404~416) were enhanced. It’s also worth noting that there were 9 potential epitopes after V367F mutation, in which the overall immunogenicity was reduced.

#### G476S

Via the mentioned predicting tool before, there are 9 changes that directly affected the effective epitope as follows (Table S7): (1) The previously likely epitopes at site of 62~75 and 459~464 were deleted. (2) The antigenicity of epitope at site of 216~221 was decreased below the threshold which cannot be an effective epitope. (3) The antigenicity of formerly ineffective epitope was elevated to be a effective epitope at site of 314~321. (4) Two epitopes (372~374+384~390) fused into a new epitope (368~390), but its antigenicity was lower than the epitope before. (5) Three alterations of epitopes improved their antigenicity with site 406~417,440~450 and 657~663. (6) The epitope at site 486~492 was reduced antigenicity slightly. Therefore, 9 effective epitopes have predicted after G476S mutation, in which the amount of epitope-changing is the most, and the overall antigenicity was dereased.

#### V483A mutation

Through the above methods for analysis, we found that 13 changes influenced significantly on B-cell potential epitope of V483A variant (Table S8). These major changes can be grouped into four categories: (1) Deletion: the site of 62~75 and 487~492. The antigenicity of epitope (210~221) was too low to be a suitable epitope. (2) Additional epitopes: the antigenicity of epitopes at 181~186,342~353,363~377and 617~628 were upregulated above threshold to become new epitopes. (3) Reduced antigenicity: the antigenicity of original epitope (405~418) was decreased generally at site of 405~413. (4) Improved antigenicity: four epitopes of 379~389, 442~447, 458~463, and 698~709 elevated their antigenicity. Therefore, there were 13 effective epitopes in total, and the overall antigenicity after mutation was advanced compared with prototype spike protein.

#### D614G mutation

D614G mutation was of higher frequency in global area (6294/10022 samples). The SARS-CoV2 strains with G614 form was more transmissible. Once it entered the new area, it could rapidly become the dominant infection form according to the epidemiological analyze [12]. Therefore, we paid a great attention to analyze the immunological characteristics of this mutation. We found total 29 B-cell linear epitopes. Among them, 12 effective epitopes were predicted. Surprisingly, only one epitope changed slightly when comparing with prototype S protein. The B-cell epitope 657-664 of prototype shorten one AA in D614G. This changed epitope 657-663 is with a slight increase in accessibility and antigenicity by 36.8% and 25.6%, suggesting that G614 mutant is more likely to bind to neutralizing antibodies and the binding efficacy is also increased compared with D614 stain. The remaining epitopes were consistent with the non-mutated S protein (Table S9). **Importantly,** three epitopes in the RBD domain (_384_PTKLNDL_390_, _405_DEVRQIAPGQTGKI_418_, and _487_NCYFPL_492_) are identical between D614 and G614 forms.

#### V615I mutation

After V615I mutation of S protein, only one change affected the potential epitope at site of 657~663 with slight increase of antigenicity (Table S10), the rest of 11 as same as prototype. Finally, 12 potent epitopes were predicted similarly to D614G variant.

#### V615F mutation

The background of V615F mutation is consistent with V615I. Through the above-mentioned forecasting tool, we found that 12 alterations directly affected B cell epitope other than V615I mutation obviously (Table S11). Specifically, the antigenicity of the four epitopes decreased at site of 140~154,209~212,441~444 and 487~497,in which the antigenicity of 209~212 and 441~444 were too low to effect. The original likely epitope of 384~390 was deleted. However, the antigenicity of four epitopes improved at site of 404~418,458~466, 656~664 and 697~709. There are three neoantigen epitopes at site 180~186,214~221 and 374~389 with strong antigenicity. Ultimately, 12 potent epitopes were predicted.

#### A831V mutation

Although A831V mutation is emerging only in Iceland as a single lineage up to now, it is found a potential fusion peptide in S2 [12],which directly affect the pathogenicity of SARS-CoV-2.After A831V mutation of S protein, 5 potential epitopes have altered (Table S12), but the remaining epitopes were consistent with those prototype. Only the antigenicity of epitope at site of 141~153 was reduced, while other 4 epitopes greatly improved their antigenicity at site 405~417, 459~465 618~625 and 657~663, in which the site (618~625) became new effective epitopes, as antigenicity increases beyond the threshold. Therefore, the S protein with A831V mutation had 13 potential epitopes, with increasing antigenicity.

### Comparison of the changes on epitopes of variants

In order to investigate the influences of the above common 9 mutations of S protein on B cell epitopes, we compared the predicted epitopes of reference and mutant S protein, analyzed the association of epitope changes among mutations and determined the influence of mutation on B cell epitopes. The detailed information of changes in each mutation is listed in the Table S13. We found that some mutations did not or slightly change B-cell epitopes, while others strongly impact the number and site of B-cell epitopes. All the major changes of B-cells comparison was summarized in Table 5. Most important finding is that the commonest mutation D614G change the B-cell epitopes of S protein slightly, only moderately increasing the accessibility and antigenicity of epitope ^657-663^. There are 12 potential epitopes in D614G mutation, nearly identical to those without mutation. In D614G and V615I mutation, their effective epitopes were also 12, in which only 1 epitope at the same site of 657~663 could affect the potential effective epitopes with slight increase of antigenicity. However, the amounts and forms of changes in epitopes were abundant in V483A, V615F, V367F and G476S mutations. Among them, the alterations of epitopes in V483A are the most significant, 13 changes were present and 13 epitopes predicted, and the antigenicity was highly improved. Changes in V615F followed by, 11 effective epitopes and 12 changes were discovered with reduced antigenicity as a whole. In addition, the change sites of some epitopes are common to several mutants. The change in epitopes 459~464 occurred in many mutations: Y145H, V367F, G476S, V483A, V615F and A831V. The epitope of 657~663 alters in Y145H, G476S, D614G, V615I, V615F and A831V. The epitope at 1154~1169 doesn’t change among 9 mutations.

## Discussion

As we know, a majority of COVID-19 vaccines at the present stage were designed to target S proteins in order to induce the neutralizing antibody [15, 16], and most vaccines entering phase 3 clinical trials are based on the early S protein [17, 18]. However, the massive replication and rapid global transmission of SARS-CoV-2 provide the virus with sufficient opportunity to mutate and evolve.

By searching the literature and database of SARS-CoV-2, we found 11 common mutations of S protein shown in Table 1, and 5 of them are concentrated on and near RBD domain. Especially, the most frequently occurring mutation D614G located at S1 and S2 junctions, where is near the furin cleavage site of the S1/S2 boundary. Walls reported that deletion of this cleavage region could influence SARS-CoV-2 S-mediated entry into host cells [8]. Hence, Korber [12] and Zhang [19] proposed that D614G mutation contributes to the spread of SARS-CoV-2, which makes G614 strain swiftly become the dominant mutant.

The mutation in S protein may affect the B–cell epitopes and lead to vaccine failure. Therefore, in order to explore the impact of mutations on antigenicity of S protein, in this study, we applied immuno-informatics tools to predict potential B-cell epitopes of prototype and variant S protein.

The reliably of prediction tools and methods were demonstrated by previous studies, for example, the epitopes prediction of MERS-CoV. Qamar reported that the linear and discontinuous epitopes were successfully predicted with the same immuno-informatics tools as we used here [20]. In other confirmed experiments, hMS-1 mAbs (monoclonal antibody) recognized some discontinuous epitopes predicted from Qamar’s research [21].

Our predicted epitopes of prototype S protein also coincide with sites of neutralizing antibodies validated in other group’s experimental studies. Cao et al have firstly shown that 7 mAbs isolated from 60 convalescent patients of COVID-19 showed both strong affinity binding to RBD and a potent neutralizing ability against SARS-CoV-2 [22]. They possessed high structural similarity with m396, previously neutralizing antibody of SARS-CoV, which can recognize epitopes (residues 408, 442, 443, 460, 475) on the RBD domain of SARS-CoV S protein [23]. Since RBD region of S protein is more prone to neutralizing antibodies [24], we checked epitopes in this 318~550 RBD region. The validations of these experiments above are consistent with our predicted B cell epitopes as follows: 5 potential linear epitopes as 384~390, 405~418, 441~448, 459~464 and 487~492 (Table 2) and discontinuous epitopes such as No.2 (Table 3) could be vaccine candidates targets.

Importantly, it’s worth exploring whether or not the mutations on S protein leads to epitope changes. Therefore, we used a group of prediction tools of B-cell epitopes to predict the prototype and variant S protein.

At present, the most notable mutation is D614G, the most popular dominant mutation among genetic variants (6294 D614G in10 022 cases, 63%) [9]. Fortunately, we demonstrate that S protein with D614G mutation has the least change of the potential effective B-cell epitope, in which only one linear epitope has slight change in the length of non-RBD region, compared with the prototype. The predicted B-cell epitopes on RBD domain are same between D614 and G614 forms. Hence, this result suggests that the effective vaccine based on prototype S protein should protect the infection of both prototype and D614G variant virus, which cover more than 90% cases estimated from data of Koyama’s study[9]. That is to say, the vaccines currently being developed could protect a large proportion of the SARS-CoV-2 infected population, including both the D614 prototype and the dominant G614 variant. Our prediction results are consistent with Weissman et al’s study [25], in which G614 Spike pseudovirions were not less susceptible to neutralization, but instead moderately more. This data indicate that D614G mutation do not change the B-cell epitopes to escape the immune recognition.

Based on the prediction results, we also found that some mutations significantly change the B-cell epitopes, for example, G476S and V483A, which perhaps impact the effect of SARS-CoV-2 vaccine with prototype S protein. The vaccination of SAES-CoV-2 vaccine may give a selection pressure for these variants.

On the other hand, we found that some mutations significantly change the B-cell epitopes. The predicted effective epitopes of D614 form are missing or down-regulated at some mutated S protein, such as V367F, G476S, V483A and V615F. These variants with more changes in potential epitopes may have significant differences in response to vaccines. If responsiveness to vaccines reduced, it will select the SARS-CoV-2 viral strains. However, this low-response epitope accounts for a small proportion of potential epitopes, so the reactive down-regulation of the vaccine is also limited and does not fundamentally alter the efficacy of vaccine. Anyhow, these variants were rare in COVID-19 population. Therefore, the upcoming vaccine may have positive effects on major SARS-CoV-2.

In conclusion, by the prediction of B-cell epitopes of the prototype and different mutant S proteins, we demonstrate that some mutations could significantly change B-cell epitopes, but others just slightly. Fortunately, the dominant variant D614G nearly do not change the B-cell epitopes of prototype S protein. This indicates that the vaccine based on prototype S protein could effectively protect human population against most of infection of SARS-CoV-2 viruses. This would be beneficial to the large-scale application of prototype vaccines of SARS-CoV-2, including in high incidence areas rich in mutant strains. We should also be of vigilance that as the application of SARS-CoV-2 vaccine, it would lead to selection pressure for S protein variants that maybe significantly change B-cell epitopes with antigen escape. International society should prepare for this situation after SARS-CoV-2 vaccination in advance.

## Materials and Methods

### Data retrieval and number of variant analysis

The primary sequence of SARS-CoV-2 S protein was retrieved from NCBI GenBank database using accession number QHO62107.1 and was used as prototype sequence or reference sequence for vaccine development in many projects [14]. Its complete genome number is NC_045512, which. The major variation sequences were available from The Global Initiative for Sharing All Influenza Data (GISAID) [26] and GenBank database. As for epidemiological statistics on the number of mutations, we selected data collected from two time points to observe the change of the amount of mutations. One data set collected until April 13, 2020 was reported by Korber B in a pre-print paper [12], while another until May 1, 2020 was from Takahiko Koyama’s research[9]. We performed the secondary classification analysis on the 10 022 sequences listed in the supplement data of Takahiko Koyama’s paper, and obtained the number of different S protein mutations in order to calculate the mutation frequency. We then selected 10 variants with a amount of mutations greater than 10 cases for further analysis.

### Conservation analysis of selected S protein sequences

S protein sequences from 10 different countries were randomly achieved from an open NCBI Genbank database. By utilizing Clustal Omega tool (Version 1.2.4) [27] and MSAViewer tool from VIPR database (https://www.viprbrc.org), multiple sequence alignment (MSA) was carried out to perceive the conservation of sequence twice [28]. MSAViewer tool could provide the visual comparison results. The aligned files were additionally applied to make phylogenetic tree via Clustal Omega Self-contained analytical tools (https://www.ebi.ac.uk/Tools).

### Structural and antigenicity analysis

First, we analyzed the secondary structure of S protein of SARS-CoV-2. The Conserved Domain Database (CDD) tool [29] in the NCBI website was used to analyze the main functional domains of S proteins and to determine the detail functional domains of S proteins with reference of Jun Lan’s study [13]. An TMHMM online tool (http://www.cbs.dtu.dk/services/TMHMM/) was used to examine the transmembrane topology of S protein [30]. The homologous modeling of S protein was carried out by using Swiss-model tool (https://swissmodel.expasy.org) to find its 3D-structure data. The confirmed 3D structure of SARS-CoV-2 S protein via electron microscopy 3.2 Å was acquired by using PDB ID: 6VYB from Protein-Data-Bank [8]. The antigenicity of S proteins was predicted by Vaxijen 2.0 in the default threshold of 0.4. This tool was developed to define antigens classification in view of the physicochemical properties of proteins rather than sequence alignment, and now is a common antigenicity assessment tool for vaccine design and available from internet (http://www.ddg-pharmfac.net/vaxijen/VaxiJen/VaxiJen.html) [31, 32].

### B-cell epitope prediction of S protein

We used the sequence from early onset SARS-CoV-2 as the wildtype or prototype and the recent variant virus as mutation strains to predict the B-cell epitopes of S protein. The S protein sequence was exclusive of the signal peptide (SP), TM and cytoplasmic region, and only the ectodomain of S protein was used for analysis. The linear and non-linear (discontinuous) epitopes of B cell were predicted by the different tools. The linear epitopes were prediced by BepiPred-2.0 server of IEDB online database [33, 34]. The threshold was set to 0.55, which represented that the sensitivity was 29%, and the specificity was 81%. Analysis result shows in a figure in which the residues with scores above the threshold predicted to be part of an epitope were colored in yellow. The effective B-cell epitopes relies on stronger antigenicity and accessibility of surface [35]. Then the better epitopes were evaluated by Vaxijen 2.0 for total antigenicity scores and Emini surface accessibility prediction tool for accessibility scores [36]. The prediction of discontinuous epitopes depended on surface accessibility and amino acid statistics and X-ray crystallography of protein epitopes [37]. We predicted the discontinuous epitopes of prototype S protein via DiscoTope 2.0 server [38] by entering PDB ID number: 6VYB. The threshold was determined at 0.5, which manifested 23% sensitivity and 90% specificity.

### Comparison of B-cell epitopes between the prototype and mutated S protein

According to the research of Korber B, et al. [12] and Takahiko Koyama [9], we selected 9 variants of the mutated S proteins for analysis of their B cell epitopes and compared each of them with epitopes from prototype sequence to determine the influence of mutation on epitopes. Finally, we summarized and listed all of major changes in a table.

### Statistical analysis

No statistical analyses were applied in this theoretical study, and the results are based on data in the published literature and publicly available databases.

## Data and code availability

All data retrieved and analyzed in the present research was obtained from the NCBI, IEDB, GISAID and PDB database and other open databases. The published literature includes all quoted or analyzed data during this study, and summarized in the figures, tables and Supplemental information.

## Author contributions

Xianlin Yuan performed the analyses and wrote the paper; Liangping Li proposed and supervised this project, wrote and revised the manuscript.

## Acknowledgments

It is grateful to Zijun Shu for his management of manuscript reference.

## Supporting information

Supporting Information can be found at the attachment.

